# Pan-arthropod analysis reveals somatic piRNAs as an ancestral defence against transposable elements

**DOI:** 10.1101/185694

**Authors:** Samuel H. Lewis, Kaycee A. Quarles, Yujing Yang, Melanie Tanguy, Lise Frézal, Stephen A. Smith, Prashant P. Sharma, Richard Cordaux, Clément Gilbert, Isabelle Giraud, David H. Collins, Phillip D. Zamore, Eric A. Miska, Peter Sarkies, Francis M. Jiggins

## Abstract

In animals, small RNA molecules termed PIWI-interacting RNAs (piRNAs) silence transposable elements (TEs), protecting the germline from genomic instability and mutation. piRNAs have been detected in the soma in a few animals, but these are believed to be specific adaptations of individual species. Here, we report that somatic piRNAs were likely present in the ancestral arthropod more than 500 million years ago. Analysis of 20 species across the arthropod phylum suggests that somatic piRNAs targeting TEs and mRNAs are common among arthropods. The presence of an RNA-dependent RNA polymerase in chelicerates (horseshoe crabs, spiders, scorpions) suggests that arthropods originally used a plant-like RNA interference mechanism to silence TEs. Our results call into question the view that the ancestral role of the piRNA pathway was to protect the germline and demonstrate that small RNA silencing pathways have been repurposed for both somatic and germline functions throughout arthropod evolution.

## Introduction

In animals, 23–31 nucleotide (nt) PIWI-interacting RNAs (piRNAs) protect the germline from double-strand DNA breaks and insertion mutagenesis by silencing transposons^1–3^. In *Drosophila*, piRNAs also function in gonadal somatic cells that support oogenesis^4,5^. Although the role of piRNAs in the germline appears to be deeply conserved across animals, they have also been reported to function outside the germline. In the mosquito *Aedes aegypti*, there are abundant non-gonadal somatic piRNAs that defend against viruses^6,7^. In other species, piRNAs are produced in specific cell lineages. For example, somatic piRNAs silence transposons in *D. melanogaster* fat body^8^ and brain^9,10^, they are important for stem cell maintenance and regeneration in the planarian *Schmidtea mediterranea*^11,12^, and they contribute to memory in the central nervous system of the mollusc *Aplesia californica*^13^.

piRNA pathway genes in *Drosophila* species evolve rapidly, likely reflecting an evolutionary arms race with TEs^14,15^. Expansion and loss of key genes in the piRNA pathway has occurred in platyhelminths^16^, nematodes^17^, and arthropods^18–20^. This gene turnover is accompanied by a wide variety of functions for piRNAs, such as sex determination in the silkworm *Bombyx mori* and epigenetic memory formation in the nematode *C. elegans*^21^. There is also considerable divergence in downstream pathways linked to piRNA silencing—for example, in *C. elegans* where piRNAs act upstream of an RNA-dependent RNA polymerase (RdRP) pathway that generates secondary siRNAs antisense to piRNA targets. Moreover, many nematode species have lost the piRNA pathway altogether, with RNAi-related mechanisms assuming the role of TE suppression^22^. These examples highlight the need for further characterisation across animals to better understand the diversity of the piRNA pathway.

To reconstruct the evolutionary history of small RNA pathways, we sampled 20 arthropod species with sequenced genomes: three chelicerates, one myriapod, one crustacean, and 15 insects. For each species, we sequenced long and small RNAs from somatic and germline adult (Supplementary Table 1). Our results highlight the rapid diversification of small RNA pathways in animals, challenging previous assumptions based on model organisms. First, we find that RdRP was an integral part of an ancestral siRNA pathway in early arthropods that has been lost in insects. Second, we demonstrate that somatic piRNAs are an ancestral trait of arthropods. Intriguingly, the somatic piRNA pathway is predominantly targeted to transposable elements, suggesting that the piRNA pathway was active in the soma of the last common ancestor of the arthropods to keep mobile genetic elements in check.

## Results & Discussion

### Extensive turnover in arthropod small RNA pathways

The duplication or loss of small RNA pathway genes can lead to the gain or loss of small RNA functions. To identify expansions of small RNA genes throughout the arthropods, we identified homologs of key small RNA pathway genes and used Bayesian phylogenetics to reconstruct the timing of duplication and loss (Fig. 1a). Small RNAs bind to Argonaute proteins and guide them to their RNA targets. siRNAs are associated with Ago2-family Argonautes, and these have been extensively duplicated across the arthropods, with an ancient duplication in the arachnid (spider and scorpion) ancestor, and lineage-specific duplications in the scorpion *Centruroides sculpturatus*, the spider *Parasteatoda tepidariorum,* the locust *Locusta migratoria*, and the beetle *Tribolium castaneum*^23^. piRNAs are associated with PIWI-family Argonautes, which have undergone similar duplications. *Piwi* has duplicated in *L. migratoria,* the centipede *Strigamia maritima*, the pea aphid *Acyrthosiphon pisum*^18^, the mosquito *Aedes aegypti* ^24^, and flies (generating *piwi* and *aubergine*^19^). All species harbour a single copy of *ago3*, which encodes the other PIWI-family Argonaute associated with piRNAs, except for *A. pisum* which has two *ago3* genes.

**Figure 1:**
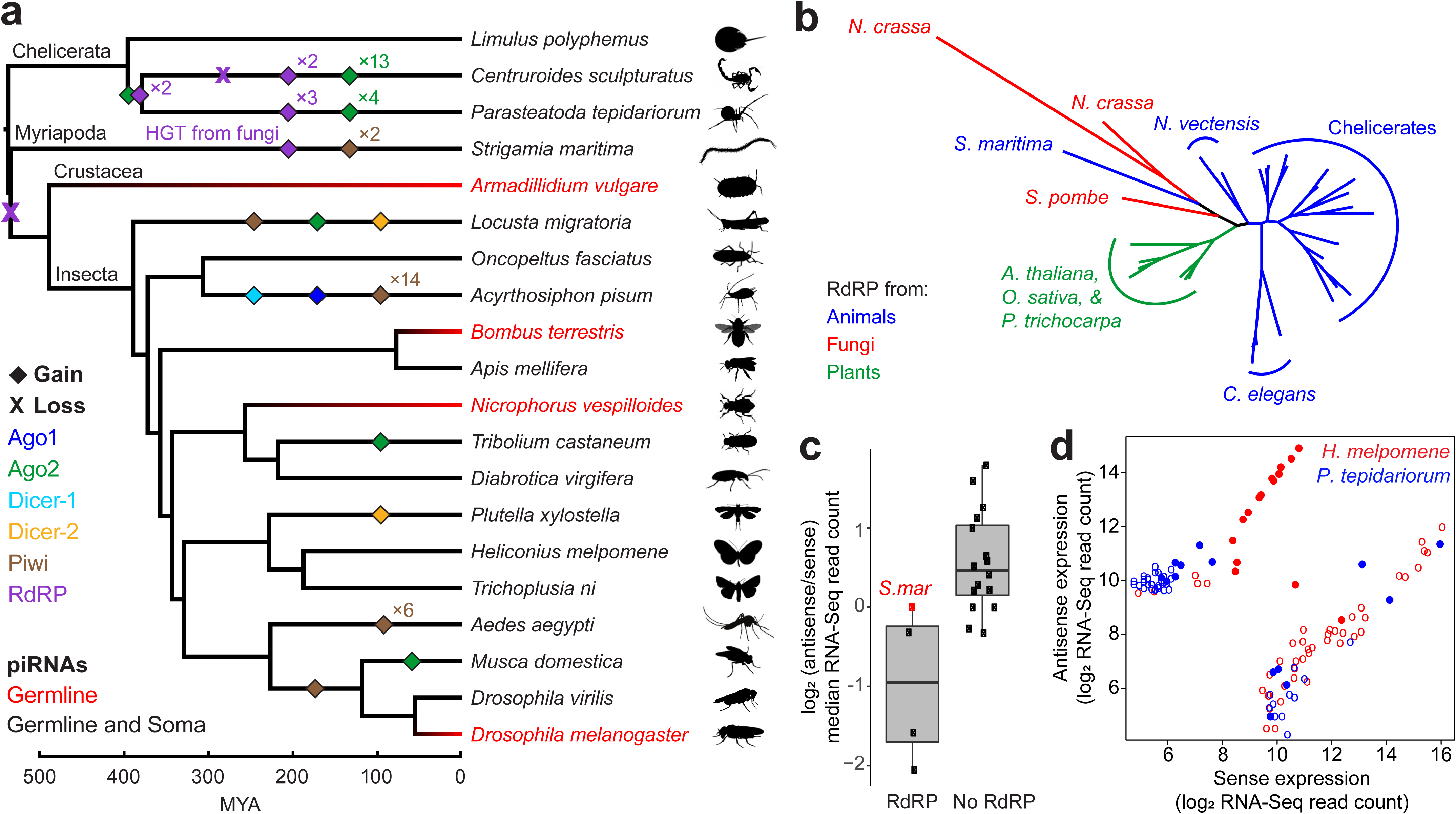
Genes in small RNA pathways evolve rapidly throughout the arthropods. **a,** The gain and loss of genes encoding the components of different sRNA pathways during arthropod evolution. Taxa with somatic piRNAs are shown in black, and the colour of the branches is a Bayesian reconstruction of whether somatic piRNAs were present. The posterior probability that the ancestral arthropod had somatic piRNAs is 0.9956. **b**, Phylogenetic analysis of RdRP genes from arthropods, other animals, plants and fungi. Note *S. maritima* is more closely related to fungal than animal RdRP (posterior probability at *N. crassa* - *S. maritima* node is 1). **c**, The antisense enrichment (measured as log_2_ (antisense/sense) median RNA-Seq read counts) for TEs that produce siRNAs. Species are classified by possession of an RdRP. Note *S. maritima* (red) lacks an animal RdRP. **d**, Counts of sense and antisense RNAseq reads of the 60 most highly expressed TEs in *H. melpomene* (no RdRP; red) and *P. tepidariorum* (six RdRPs; blue). Among these, the 15 TEs in each species that generate the most siRNAs are shown as filled circles while the remainder are open circles. In *H. melpomene* siRNAs are associated with antisense transcription.

RdRPs amplify an siRNA signal by generating double-stranded RNA (dsRNA) from single-stranded RNA^25^, but *Drosophila* and other insects lack RdRP genes. RdRP is present in some ticks^26^, and similarly, we identified RdRP genes across the chelicerates, frequently in multiple copies (Fig. 1a). In each species, one or more RdRPs are expressed in at least one tissue (Supplementary Fig. 1). We also identified an RdRP in the centipede *S. maritima*; however, phylogenetic analysis provides strong evidence that this is not an orthologue of the ancestral arthropod RdRP, but is more closely related to RdRP from fungi (*Neurospora crassa* and *Schizosaccharomyces pombe*; Fig. 1b). In contrast, the chelicerate RdRP is most closely related to other animal RdRPs. Given that RdRPs are present in nematodes and *Nematostella vectensis*, the most parsimonious explanation is that RdRP was present in the common ancestor of arthropods and has been retained in the chelicerates. It was then lost in all other arthropods ∼500 MYA, and subsequently regained by *S. maritima* by horizontal gene transfer from a fungus (Figs. 1a,b).

The RdRPs expressed in the chelicerates and *S. maritima* may generate dsRNA precursors which can then be processed by Dicer to generate siRNAs, similar to RdRPs in basal nematodes^22^, while species lacking an RdRP would require bidirectional transcription by RNA polymerase II to generate dsRNA. To test this idea, we sequenced long RNA (RNA-Seq) and small RNA from all species (Supplementary Table 1). Within each species, we identified TEs that were expressed and targeted by siRNAs, and estimated the difference between their sense and antisense expression. Compared to species lacking RdRPs, we find that species with RdRPs have less antisense transcription of these TEs (Mann-Whitney *U* test, animal RdRP versus no RdRP: *p* = 0.0381; Fig. 1c). This pattern is also apparent when comparing antisense transcription and siRNA production across the 15 most highly-expressed TEs within a single species. For example, in *H. melpomene*, which does not have an RdRP, there is a significant positive correlation between the proportion of antisense transcripts and siRNA production (Spearman rank correlation ρ = 0.52, *p* = 2×10^−5^). Furthermore, none of the TEs with low antisense transcription are among the top siRNA targets (Fig. 1d). These results suggest that *H. melpomene* requires bidirectional transcription to generate siRNAs. In contrast, in *P. tepidariorum* (six RdRPs) there is no correlation between the proportion of antisense transcripts and siRNA production (Spearman rank correlation ρ = 0.09, *p* = 0.512), and several TEs with very few antisense transcripts generate abundant siRNAs (Fig. 1d). Together, our results suggest that chelicerates are less dependent on bidirectional transcription to provide the precursors for siRNA production, and may use RdRP to generate dsRNA from TEs, similar to plants and some nematodes. However, we note that the antisense enrichment for siRNA targets in *S. maritima* is more similar to species lacking an RdRP, making it unclear whether its horizontally-transferred RdRP acts in this way.

### Germline piRNAs are found across arthropods

Current evidence supports the view that the piRNA pathway is a germline-specific defence against transposon mobilization. As expected, we found piRNAs derived from the genome in the female germline of all 20 arthropod species (Supplementary Table 1, Supplementary Fig. 2), consistent with deep conservation of this function from the last common ancestor of mammals and arthropods. Germline piRNAs target TEs in a wide variety of animals, including nematodes, fish, birds, and mammals, as was the case in all our species (Supplementary Fig. 3); moreover, TE abundance and piRNA abundance were positively correlated as previously found in *D. melanogaster* (Supplementary Fig. 4). In 10 species, we also sampled the male germline. Male germline piRNAs were found in all species except the bumblebee *Bombus terrestris*, which lacked detectable piRNAs in both testis and mature sperm-containing vas deferens, even when using a protocol that specifically enriches for piRNAs by depleting miRNAs^9^ (Fig. 2a; Supplementary Figs. 5 and 6). In contrast, piRNAs were abundant in *B. terrestris* ovary (Fig. 2b; Supplementary Fig. 2). Moreover, mRNAs encoding the core piRNA pathway proteins Piwi and Vasa were 10-fold less abundant in testis compared to ovary (Supplementary Fig. 7), suggesting that the piRNA pathway is not active in the *B. terrestris* male germline. To our knowledge, this is the first report of sex-specific absence of piRNAs in the germline, and suggests that other processes may have taken on the function of TE suppression in *B. terrestris* males. Male bumblebees are haploid and produce sperm by mitosis rather than meiosis^27^, unlike males from the other eight species analysed. However, in the testis of the haplodiploid honey bee *Apis mellifera* piRNAs are detectable by their characteristic Ping-Pong signature, albeit at low levels (Supplementary Figs. 5 and 8).

**Figure 2:**
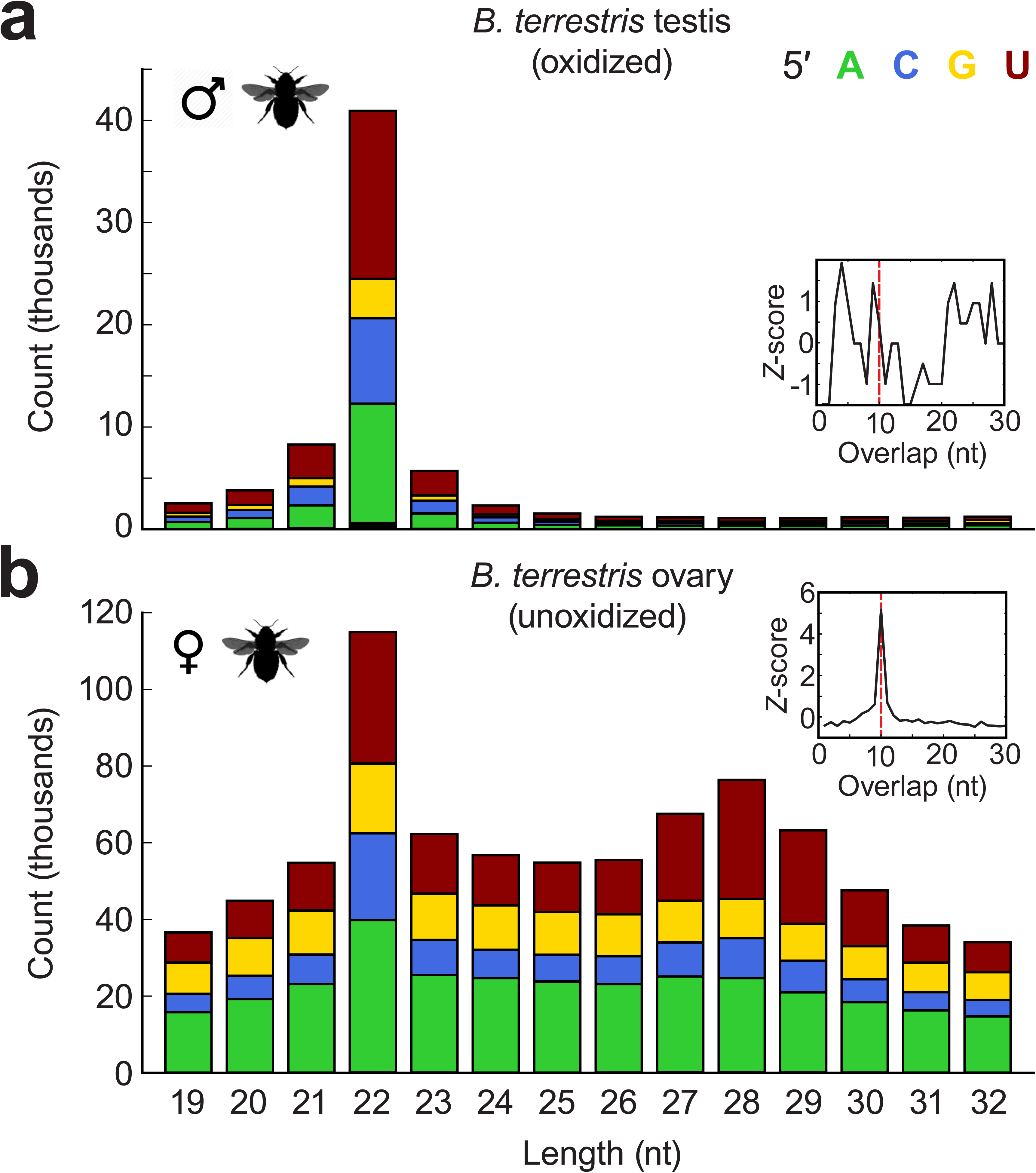
piRNAs are absent in *B. terrestris* male germline. The size and 5′ nucleotide of sRNAs from testis (**a**) and ovary (**b**). Plots show unique reads that map to the genome (where the same sequence occurred more than once, all but one read was eliminated). The inset shows the overlap between sense and antisense 25-29nt sRNAs.

**Figure 3:**
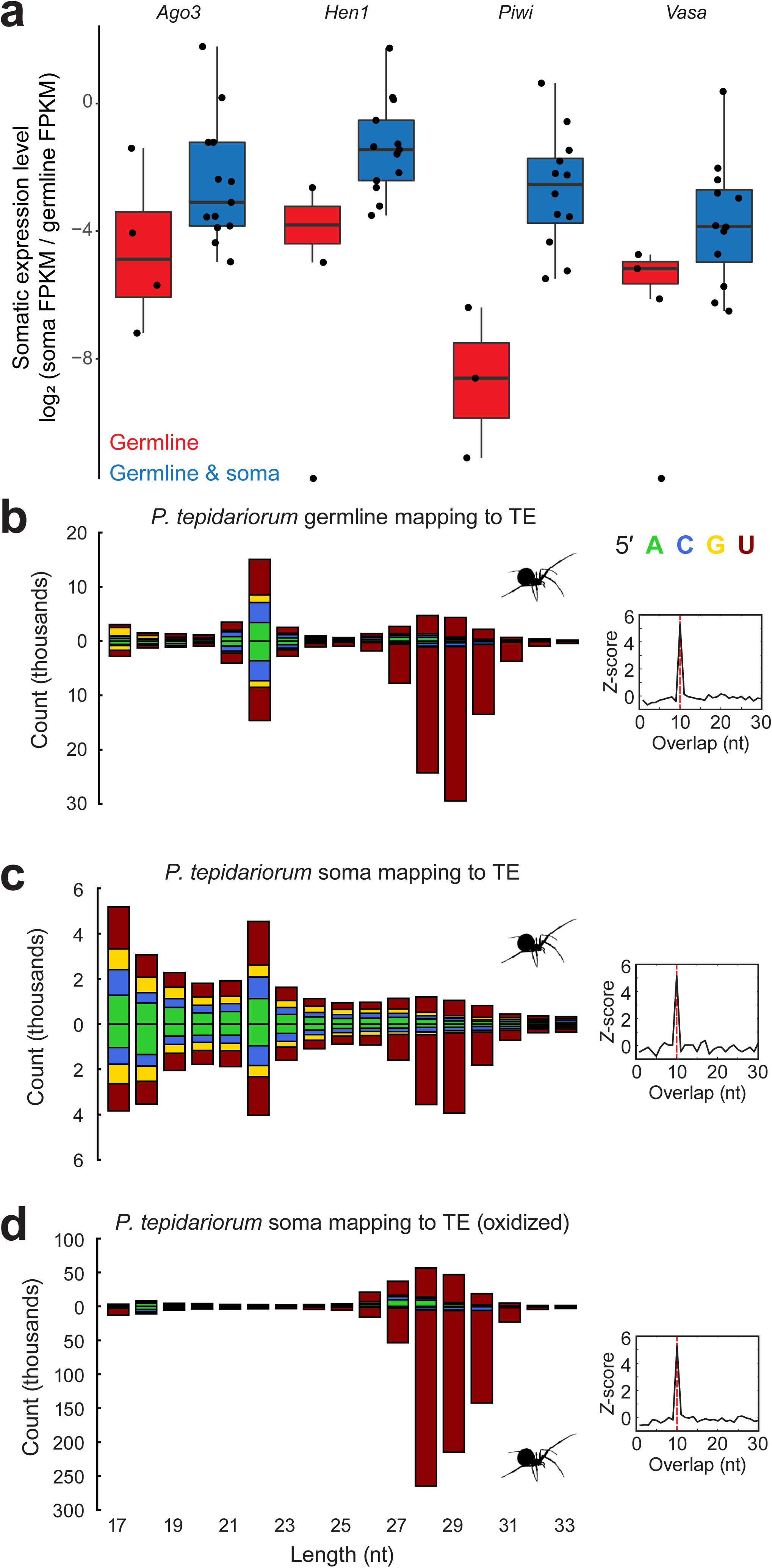
Somatic piRNAs are widespread, and target TEs throughout the arthropods. **a,** Genes in the piRNA pathway have higher somatic expression in species with somatic piRNAs. For species with multiple copies of a gene, the mean scaled somatic expression level of each duplicate is displayed. The box shows the median and interquartile range (IQR), and the bottom and top whiskers show the range of points no further than 1.5×IQR away from the first and third quartiles respectively. **b-d,** The size and 5′ nucleotide of sRNAs mapping to the TEs from *P. tepidariorum,* showing 10 bp overlap between sense and antisense 25–29nt sRNAs. piRNAs targeting TEs are evident in the germline (**b**) and soma (**c**), and these somatic piRNAs are resistant to sodium periodate oxidation, indicating that they are 3′ methylated (**d**). Plots show all reads that map to the TEs.

**Figure 4:**
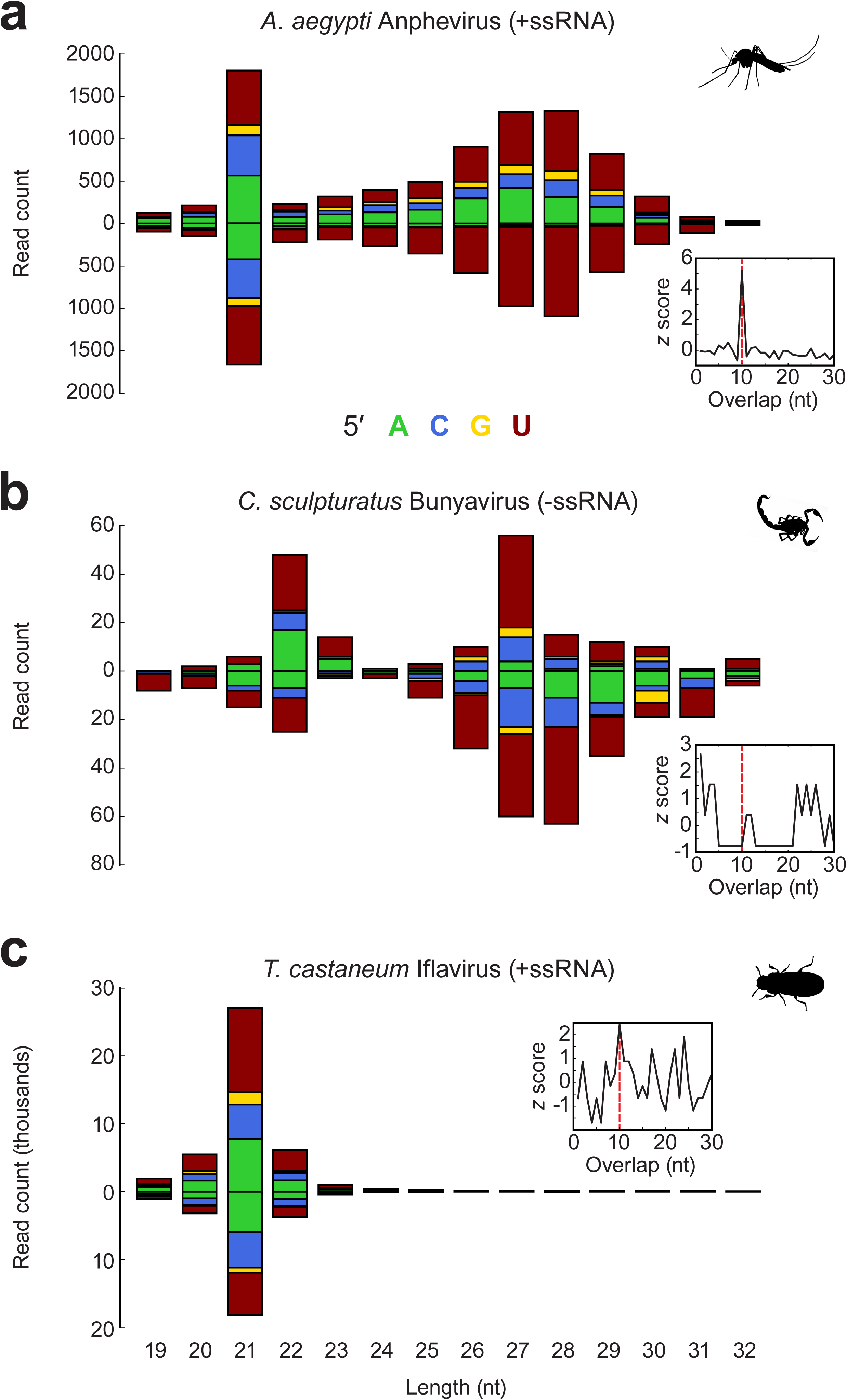
Virally-derived sRNAs in three arthropod species. The size and 5′ nucleotide of sRNAs mapping to viral transcripts and genomes reconstructed from RNA-Seq data. Virally-derived piRNAs are evident in *A. aegypti* (**a**) and *C. sculpturatus* (**b**), and virally-derived siRNAs are found in *T. castaneum* (**c**). Only *A. aegypti* shows the 10 bp overlap between sense and antisense 25–29 nt sRNAs that is diagnostic of Ping-Pong amplification (insets). Reads derived from the sense strand are shown above zero, antisense reads below.

**Figure 5:**
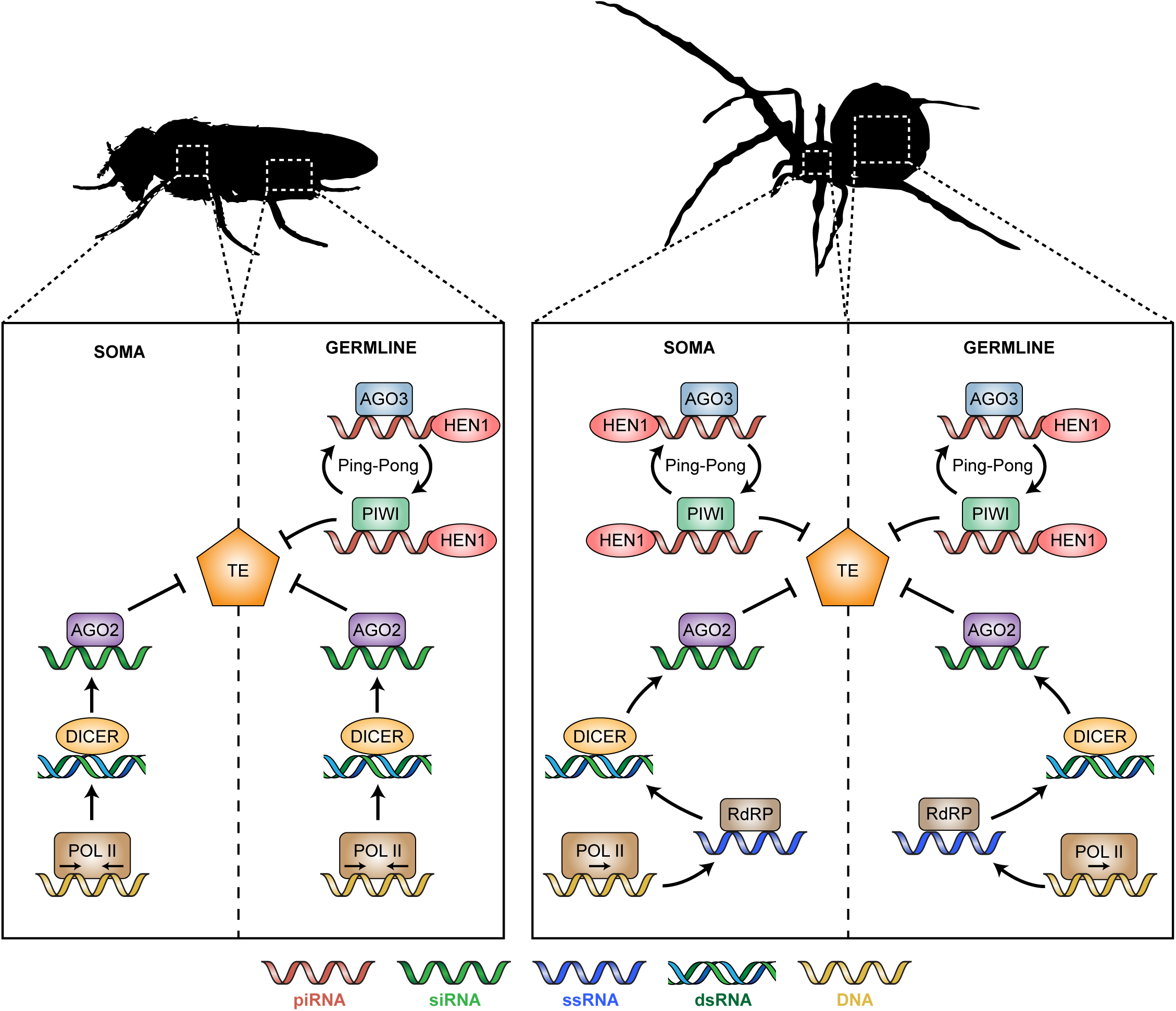
A model of the divergent sRNA pathways silencing TEs in different arthropods. Our data suggest that the mechanisms of sRNA pathways have diverged in two key areas. In some lineages, the piRNA pathway is restricted to the germline (e.g., flies), whereas in most others it is active in the soma and the germline (e.g., spiders). Additionally, in some lineages (e.g., spiders), RdRP may synthesize dsRNA from transcripts produced by RNA polymerase II, amplifying the siRNA response.

### Somatic piRNAs are widespread across arthropods

Among the 20 arthropods we surveyed, somatic piRNAs were readily detected in 16 species: three chelicerates (*L. polyphemus*, *C. sculpturatus*, and *P. tepidariorum*), the myriapod *S. maritima*, and 12 insect species (Figs. 1a and 3c,d; Supplementary Fig. 9). We did not detect piRNAs in the somatic tissues of the crustacean *Armadillidium vulgare* or the insects *N. vespilloides*, *B. terrestris*, and *D. melanogaster* (Supplementary Fig. 9). Although somatic piRNAs have been detected previously in *D. melanogaster* heads^9,10^, we detected no piRNAs in *D. melanogaster* thorax. Somatic expression of the piRNA pathway genes *vasa, ago3*, *Hen1*, and *Piwi* was strongly associated with the presence of somatic piRNAs (Fig. 3a). We conclude that an active somatic piRNA pathway is widespread throughout the arthropods.

The phylogenetic distribution of somatic piRNAs suggests that they were either ancestral to all arthropods or have been independently gained in different lineages. To distinguish between these possibilities, we used ancestral state reconstruction to infer the presence or absence of somatic piRNAs on the internal branches of the arthropod phylogeny. Our results indicate that somatic piRNAs are ancestral to all arthropods (posterior probability = 0.9956), and have been independently lost at least four times (Fig. 1a).

### Functions of somatic piRNAs

In all but one species with somatic piRNAs, at least 2% of piRNAs mapped to TEs (Fig. 3c, Supplementary Fig. 3), suggesting that their anti-transposon role is conserved in the soma. The exception to this pattern was *O. fasciatus*, where only 0.009% of somatic and 0.074% of germline piRNAs were derived from annotated TEs. Moreover, somatic piRNAs from all species displayed the hallmark features of piRNA biogenesis and amplification: a 5′ uracil bias, 5′ ten-nucleotide complementarity between piRNAs from opposite genomic strands (“Ping-Pong” signature), and resistance to oxidation by sodium periodate, consistent with their bearing a 2′-*O*-methyl modification at their 3′ ends (e.g., Fig. 3d). Given the ubiquity of TE-derived somatic piRNAs, we wondered whether there was a relationship between the TE content of a species’ genome and the presence of somatic piRNAs. However, although species with somatic piRNAs tend to have a higher TE content, this difference is not significant (*p* = 0.18, Supplementary Fig. 10).

In *Drosophila*, piRNAs derived from protein-coding genes are thought to play a role in regulating gene expression^28^. Somatic piRNAs derived from protein-coding sequences and untranslated regions (UTRs) were present in all species possessing somatic piRNAs except *A. mellifera*, *D. virilis* and *M. domestica*, which lack both a distinct peak of 25-29nt sRNAs and a Ping-Pong signature (Supplementary Fig. 3, Supplementary Fig. 11). When scaled to the genome content of each feature, there is no consistent difference in the abundance of piRNAs from protein-coding sequence and UTRs (Supplementary Fig. 12), suggesting that somatic piRNAs target genes across the entire length of the transcript, rather than just UTRs.

In the mosquito *A. aegypti*, somatic piRNAs target viruses^6,7^. To test whether somatic piRNAs derive from viruses in other species, we reconstructed partial viral genomes from each species using somatic RNA-Seq data, then mapped small RNAs from these tissues to these viral contigs. In *A. aegypti*, we recovered the partial genome of a positive-sense, single-stranded RNA virus that was targeted by both siRNAs (21 nt) and 5′ U-biased, 25–30 nt piRNAs bearing the signature of Ping-Pong amplification (Fig. 4a). These data recapitulate previous results showing that both the siRNA and piRNA pathways mount an antiviral response in *A. aegypti*^6^, and thus validate our approach. In eight additional species, we could similarly reconstruct viruses that generated antiviral siRNAs (Fig. 4c, Supplementary Fig. 13). Four of these species also produced 25–30 nt, 5′ U-biased RNAs derived from viruses including negative- and positive-sense RNA viruses and DNA viruses (Fig. 4b, Supplementary Fig. 13). There was no evidence of Ping-Pong amplification of viral piRNAs in any of these species—in *C. sculpturatus* somatic piRNAs were of low abundance (Fig. 4b), and in *T. castaneum*, *D. virgifera* and *P. xylostella* piRNAs mapped to only one strand (Supplementary Fig. 13), a feature reminiscent of the somatic piRNAs present in *Drosophila* follicle cells^4,5^. Despite removing sequencing reads that map to the reference genome, we cannot exclude the possibility that these piRNAs come from viruses integrated in the host genome^29^. Together these results suggest that although some viruses may be targeted by somatic piRNAs, siRNAs likely remain the primary antiviral defence against most viruses across the arthropods.

## Conclusions

The rapid evolution of small RNA pathways makes inferences drawn from detailed studies of individual model organisms misleading^22^. Our results suggest that the best studied arthropods, concentrated in a small region of the phylogenetic tree, are not representative of the entire phylum (Fig. 5). First, ancestral arthropods likely used an RdRP to generate siRNAs from transposable elements. RdRPs likely expand the range of substrates that can generate siRNAs, because these RNA-copying enzymes provide an alternative to the generation of dsRNA precursors by RNA polymerase II. Second, and more surprising, somatic piRNAs are ubiquitous across arthropods, where they target transposable elements and mRNAs. The rapid and dynamic evolution of somatic and germline piRNA pathways across the arthropods highlights the need for a deeper examination of the origins and adaptations of the piRNA pathway in other phyla.

## Methods

### Tissue dissection

To sample germline tissue from each species, we dissected the female germline of all 20 arthropods (ovary and accessory tissue). For *Limulus polyphemus*, *Centruroides sculpturatus*, *Parasteatoda tepidariorum*, *Armadillidium vulgare, Locusta migratoria*, *Bombus terrestris*, *Apis mellifera*, *Nicrophorus vespilloides*, *Heliconius melpomene* and *Trichoplusia ni*, we also dissected the male germline (testes, vas deferens, and accessory tissue). We were unable to isolate sufficient germline tissue for *Strigamia maritima*.

To isolate somatic tissue, we used different dissection approaches depending on the anatomy of the species. In each case, we minimized the risk of germline contamination by selecting tissue from either a body region that was separate (e.g., thorax) or physically distant from the germline. For insects, thorax served as a representative somatic tissue. For *Oncopeltus fasciatus*, *Acyrthosiphon pisum*, *Apis mellifera*, *Tribolium castaneum*, *Diabrotica virgifera*, *Plutella xylostella*, *Aedes aegpyti, Musca domestica* and *Drosophila melanogaster* we used female thorax; for *Locusta migratoria*, *Bombus terrestris*, *Nicrophorus vespilloides*, *Heliconius melpomene,* and *Trichoplusia ni* we used female and male thorax separately. For non-insect species, we took mixed tissue from either the mesosoma (*Parasteatoda tepidariorum*), prosoma (*Centruroides sculpturatus*), pereon and pleon (*Armadillidium vulgare*) or muscle, heart, and liver (*Limulus polyphemus*). For these non-insect species, we isolated somatic tissue from males and females separately. For *Strigamia maritima*, we pooled female and male fat body.

### RNA extraction and library preparation: Protocol 1

For *Limulus polyphemus*, *Centruroides sculpturatus*, *Parasteatoda tepidariorum*, *Strigamia maritima*, *Armadillidium vulgare, Locusta migratoria*, *Bombus terrestris*, *Nicrophorus vespilloides* and *Heliconius melpomene* we extracted total RNA and constructed sequencing libraries using Protocol 1. Following dissection, each sample was homogenized in Trizol (Invitrogen, Carlsbad, CA, USA) and stored at –80°C. RNA from each sample was extracted with isopropanol/chloroform (2.5:1), and RNA integrity was checked using the Bioanalyzer RNA Nano kit (Agilent, Santa Clara, CA, USA).

For small RNA sequencing, each sample was initially spiked with *C. elegans* RNA (N2 strain) at 1/10^th^ mass of the input RNA (e.g., 0.1 μg *C. elegans* RNA with 1 μg sample RNA). This allowed us to quantify the efficiency of sRNA library production. To sequence all small RNAs in a 5′-independent manner, we removed 5′ triphosphates by treating each sample with 5′ polyphosphatase (Epicentre/Illumina, Madison, WI, USA) for 30 min. We used the TruSeq Small RNA Library Preparation Kit (Illumina, San Diego, CA, USA) according to the manufacturer’s instructions to produce libraries from total RNA. We sequenced each sRNA library on a HiSeq 1500 (Illumina) to generate 36 nt single-end reads.

piRNAs are typically 2′-*O*-methylated at their 3′ ends, which makes them resistant to sodium periodate oxidation. To test for the presence of modified 3′ ends, we resuspended RNA in 5× borate buffer, treated with sodium periodate (25 mM f.c., e.g., 5 μl 200 mM sodium periodate in 40 μl reaction) for 10 min, recovered the treated RNA by ethanol precipitation^30^ and constructed and sequenced libraries as above.

For transcriptome and virus RNA-Seq, each sample was initially spiked with *C. elegans* RNA (N2 strain) at 1/10^th^ mass of the input RNA. To remove ribosomal RNA, we treated each sample with the Ribo-Zero rRNA Removal Kit (Human/Mouse/Rat; Illumina) according to manufacturer’s instructions, then prepared strand-specific RNA-Seq libraries using the NEBNext Ultra Directional RNA Library Prep kit (New England Biolabs, Ipswich, MA, USA), with the optional User Enzyme step to selectively degrade the 2^nd^ strand before PCR amplification. RNA-Seq libraries were sequenced on a HiSeq 4000 to generate 150 nt paired-end reads (*C. sculpturatus* and *S. maritima*), or a HiSeq 2500 to generate 125 nt paired-end reads (all other species).

### RNA extraction and library preparation: Protocol 2

For *Oncopeltus fasciatus*, *Acyrthosiphon pisum*, *Apis mellifera*, *Tribolium castaneum*, *Diabrotica virgifera*, *Plutella xylostella*, *Trichoplusia ni*, *Aedes aegpyti*, *Musca domestica, Drosophila virilis* and *Drosophila melanogaster* we extracted total RNA and constructed sequencing libraries using Protocol 2. Following dissection, we washed each sample in PBS, proceeded directly to RNA extraction using the *mir*Vana miRNA Isolation kit (Ambion, Life Technologies, CA, USA) according to the manufacturer’s protocol, and precipitated RNA with ethanol. We prepared RNA-Seq libraries for each sample from 5 μg total RNA as described^31^, after first depleting rRNA using the Ribo-Zero rRNA Removal Kit (Human/Mouse/Rat; Illumina). We sequenced each library on a NextSeq 500 (Illumina) to generate 79 nt paired-end reads.

Small RNA sequencing libraries were generated as described^32^. First, we purified 16–35 nt RNA from 10–20 μg total RNA by 15% denaturing urea-polyacrylamide gel electrophoresis. Half of each sample was then treated with sodium periodate (above). We then ligated 3′ pre-adenylated adapter to treated or untreated RNA using homemade, truncated mutant K227Q T4 RNA ligase 2 (amino acids 1–249) and purified the 3′-ligated product by 15% denaturing urea-polyacrylamide gel electrophoresis. To exclude 2S rRNA from sequencing libraries, 2S blocker oligo^33^ was added to all samples before the 5′-adapter was appended using T4 RNA ligase (Ambion). cDNA was synthesized using AMV reverse transcriptase (New England Biolabs) and the reverse transcription primer 5′-CCTTGGCACCCGAGAATTCCA-3′. The small RNA library was amplified using AccuPrime Pfx DNA polymerase (ThermoFisher, USA) and forward (5′-AATGATACGGCGACCACCGAGATCTACACGTTCAGAGTTCTACAGTCCGA-3′) and barcoded reverse (5′-CAAGCAGAAGACGGCATACGAGAT-barcode(N6)-GTGACTGGAGTTCCTTGGCACCCGAGAATTCCA-3′) primers, purified from a 2% agarose gel, and sequenced on a NextSeq 500 to generate 50 nt single-end reads.

## Bioinformatics analysis

### Gene family evolution

To reconstruct duplications and losses of sRNA pathway components, we searched for homologs of *Ago1, Ago2, Ago3, Piwi, Dcr1, Dcr2, Drosha, Hen1 and Vasa*. For each species, we took the annotated protein set and used DIAMOND^34^ to perform reciprocal all-versus-all BLASTp searches against all proteins in *D. melanogaster*, and retained only the top hit in each case. Accession numbers for the genome assemblies and annotated protein sets are detailed in Supplementary Table 2. To find homologs of RdRP, which is absent from *D. melanogaster*, we took the annotated protein set for each species and used DIAMOND to perform BLASTp searches against the RdRP from *Ixodes scapularis* (ISCW018089). For proteins in the Argonaute and Dicer families, we identified domains in hits using InterProScan5^35^ with the Pfam database, and retained only those hits containing at least one of the conserved domains in these families (PAZ and Piwi for the Argonaute family, PAZ, Dicer, Ribonuclease and Helicase for the Dicer family). For each protein, partial BLAST hits were manually curated into complete proteins if the partial hits were located adjacent to each other on the same scaffold or contig. To establish the evolutionary relationships between homologs, we aligned each set of homologs as amino acid sequences using MAFFT^36^ with default settings, screened out poorly aligned regions using Gblocks^37^ with the least stringent settings, and inferred a gene tree using the Bayesian approach implemented in MrBayes v3.2.6^38^. We specified a GTR substitution model with gamma-distributed rate variation and a proportion of invariable sites. We ran the analysis for 10 million generations, sampling from the posterior every 1000 generations.

### Transposable element annotation

To annotate transposable elements (TEs) in each genome, we used RepeatMasker v4.0.6^39^ with the “Metazoa” library to identify homologs to any previously-identified metazoan TEs. In addition, we used RepeatModeler v1.0.8^40^ to generate a *de novo* Hidden Markov Model for TEs in each genome, and ran RepeatMasker using this HMM to identify TEs without sufficient homology to previously-identified metazoan TEs. We combined these two annotations to generate a single, comprehensive TE annotation file for each species. We then screened out all annotations <100 nt long. The source code for this analysis is accessible on GitHub (https://github.com/SamuelHLewis/TEAnnotator), and the TE annotation files are available from the Cambridge Data Archive (https://doi.org/10.17863/CAM.10266).

### Virus identification and genome assembly

To identify viruses, we first mapped RNA-Seq reads to the genome of the host species to exclude genome-derived transcripts, thus filtering out endogenous viral elements present in the reference genome. We then used Trinity^41^ with default settings to generate a *de novo* assembly of the remaining RNA-Seq data for each tissue, and extracted the protein sequence corresponding to the longest open reading frame for each contig with TransDecoder (https://transdecoder.github.io/), excluding all contigs shorter than 100nt. To identify contigs that were potentially of viral origin, we used DIAMOND to perform BLASTp searches against all viral proteins in NCBI (ftp.ncbi.nih.gov/refseq/release/viral/viral.1.protein.faa.gz and ftp.ncbi.nih.gov/refseq/release/viral/viral.2.protein.faa.gz, downloaded 19/10/16). To screen out false-positive hits from those contigs with similarity to a viral protein, we used DIAMOND to perform BLASTp searches against the NCBI non-redundant (nr) database (downloaded 19/10/16) and retained only those contigs which still had a virus as their top hit. The source code for this analysis is accessible on GitHub (https://github.com/SamuelHLewis/VirusFinder), and the viral contigs are available from GenBank (accession codes MG012486-MG012488).

### Small RNA analysis

To characterize sRNAs derived from the genome in each tissue of each species, we first used the FASTX Toolkit (http://hannonlab.cshl.edu/fastx_toolkit/) to screen out small RNA reads with >10% positions with a Qphred score <20 and cutadapt^42^ to trim adapter sequences from reads. We then mapped small RNAs to the genome using Bowtie2 v2.2.6^43^ in “–fast” mode, which reports the best alignment for reads mapping to multiple locations, or a randomly-chosen location if there are multiple equally-good alignments. We quantified the length distribution, base composition, and strand distribution of sRNAs mapping to the genome using a custom Python script (accessible on GitHub https://github.com/SamuelHLewis/sRNAplot), considering unique sRNA sequences only.

To characterize sRNAs targeting TEs, we used BEDTools getfasta^44^ to extract TE sequences from the genome in a strand-specific manner (according to the TE annotation for each genome, above), mapped sRNAs as detailed above, and quantified their characteristics using the same custom Python script (https://github.com/SamuelHLewis/sRNAplot), this time considering all sRNA sequences. To characterize sRNAs targeting viruses, we first screened out genome-derived sRNAs by mapping sRNAs to the genome and retaining unmapped reads. We then used the same mapping procedure as detailed above, applied to each virus separately.

To characterize small RNAs mapping to UTRs in each species (except *D. virgifera*, *D. virilis* and *D. melanogaster*), we extracted 200 nt upstream (5′ UTR) or downstream (3′ UTR) of each gene model. To ensure that these UTR sequences did not overlap with TEs, we masked any sequence that we had annotated as a TE using RepeatMasker (see above). We then screened out TE-derived sRNAs by mapping sRNAs to the TE annotations and retaining unmapped reads. These were mapped to our UTR annotations as detailed for TEs (above). For *D. melanogaster* and *D. virilis* we employed the same method but used the curated set of 5′ and 3′ UTRs from genomes r6.15 and r1.06 respectively. We excluded *D. virgifera* from this analysis as gene models have not been predicted for its genome.

For each species, we defined the presence of UTR-derived piRNAs based on the presence of >200 unique 25-29nt sequences with a 5′ U nucleotide bias. For species with somatic piRNAs, we used oxidized sRNA data to assay the presence or absence of somatic UTR-derived piRNAs. We excluded *D. virgifera* from this analysis because of a lack of annotated gene models.

To test whether piRNAs show evidence of ping-pong amplification, we calculated whether sense and antisense 25–29nt reads tended to overlap by 10 nt using the *z*-score method of Zhang et al^45–47^.

### Gene expression analysis

To quantify the expression of genes in small RNA pathways in each tissue, we first used Trim Galore (https://github.com/FelixKrueger/TrimGalore) with default settings to trim adapters and low-quality ends from each RNA-Seq mate pair. We then mapped these reads to the genome using Tophat2 v2.1.1^48^ with default settings in “–library-type fr-firststrand” mode. To calculate FPKM values for each gene we used DESeq2^49^, specifying strand-specific counts and summing counts for each gene by all exons. We excluded *D. virgifera* from this analysis because a genome annotation file is unavailable. The source code for this analysis is accessible on GitHub (https://github.com/SamuelHLewis/GeneExpression).

### Species tree reconstruction

To provide a timescale for the evolution of arthropod sRNA pathways, we combined published phylogenies of insects^50^ and arthropods^51^ with our own estimates of divergence dates and branch lengths. We first gathered homologs of 163 proteins that are present as 1:1:1 orthologues in each of our focal species. We then generated a concatenated alignment of these proteins using MAFTT^36^ with default settings, and screened out poorly-aligned regions with Gblocks^37^ in least stringent mode. We used this alignment to carry out Bayesian phylogenetic analysis as implemented in BEAST^52^, to infer branch lengths for the phylogeny of our sample species. We specified a birth-death speciation process, a strict molecular clock, gamma distributed rate variation with no invariant sites, and fixed the topology and set prior distributions on key internal node dates (Arthropoda = 568 ± 29, Insecta-Crustacea = 555 ± 33, Insecta = 386 ± 27, Hymenoptera-Coleoptera-Lepidoptera-Diptera = 345 ± 27, Coleoptera-Lepidoptera-Diptera=327±26, Lepidoptera-Diptera = 290± 46, Diptera = 158 ± 51) based on a previous large-scale phylogenetic analysis of arthropods^50^. We ran the analysis for 1.5 million generations, and generated a maximum clade credibility tree with TreeAnnotator^52^.

### TE content analysis

To compare the TE content of species with and without somatic piRNAs, we used the TE annotations derived from RepeatModeler (above) to calculate the TE content of each genome as a proportion of the entire genome size. We then tested for a difference in TE content between species with and without somatic piRNAs using a phylogenetic general linear mixed model to account for non-independence due to the phylogenetic relationships. The model was implemented using a Bayesian approach in the R package MCMCglmm^53^ based on the time-scaled species phylogeny (see above). The source code for this analysis is accessible on GitHub (https://github.com/SamuelHLewis/TEContent).

### RdRP signature

In species with an RdRP, siRNAs can be produced from loci that are transcribed from just the sense strand, as the RdRP synthesizes the complementary strand, whereas in species that lack an RdRP, siRNAs can only be produced from loci that have both sense and antisense transcription. To test the association between siRNA production and antisense transcription in each species, we first used Trimmomatic^54^ to extract sRNAs corresponding to the median siRNA length in that species. We then used Bowtie v2.2.6^43^ in “–fast” mode to map siRNAs and RNA-Seq reads to TE sequences in each genome, and generated strand-specific counts of siRNAs and RNA-Seq reads for each TE using BEDTools coverage^44^. We then calculated the enrichment of antisense expression [log2(antisense RNA-Seq reads) - log2(sense RNA-Seq reads)] at TEs with >5 RNA-Seq reads per million and >100 siRNAs per million sRNA reads in species with and without RdRP (Fig. 1c), and tested for a difference in enrichment between species with and without RdRP (excluding *S. maritima*) using a Wilcoxon unpaired test. We also plotted the 60 most highly expressed TEs for *H. melpomene* and *P. tepidariorum* and highlighted which of these loci were among the top 15 siRNA-producing TEs (Fig. 1d). The source code for this analysis is accessible on GitHub (https://github.com/SamuelHLewis/RdRP).

## Data Availability

Sequence data that support the findings of this study have been deposited in the NCBI Short Read Archive under the BioProject accession code PRJNA386859. Length distributions of TE-mapping small RNAs and raw data used to plot Figures 1c, 1d & 3a and Supplementary Figures 1, 7 & 10 are available on the Cambridge Data Repository (https://doi.org/10.17863/CAM.10266).

## Code Availability

Source code used in this study is accessible on GitHub (https://github.com/SamuelHLewis), please see Methods for details of source code used in each analysis.

## Acknowledgements

We thank A. McGregor, D. Leite, M. Akam, R. Jenner, R. Kilner, A. Duarte, C. Jiggins, R. Wallbank, A. Bourke, T. Dalmay, N. Moran, K. Warchol, R. Callahan, G. Farley, and T. Livdahl for providing arthropods. This research was supported by a Leverhulme Research Project Grant (RPG-2016-210 to F.M.J., E.A.M. and P.S.), a European Research Council grant (281668 DrosophilaInfection to F.M.J.), a Medical Research Council grant (MRC MC-A652-5PZ80 to P.S.) an Imperial College Research Fellowship (to P.S.), Cancer Research UK (C13474/A18583, C6946/A14492 to E.A.M.), the Wellcome Trust (104640/Z/14/Z, 092096/Z/10/Z to E.A.M.), and an NIH R37 grant (GM62862 to P.D.Z.).

### Author contributions

S.H.L. and K.A.Q. performed the experiments with assistance from Y.Y., M.T., L.F., S.A.S., P.P.S., R.C., C.G., I.G., D.H.C.; S.H.L., K.A.Q. & P.S. carried out computational analysis; P.D.Z., E.A.M., P.S. & F.M.J. supervised the project; S.H.L., K.A.Q., P.D.Z., E.A.M., P.S. & F.M.J. wrote the manuscript.

### Competing financial interests

The authors declare no competing financial interests.

